# Macrophage-derived developmental endothelial locus 1 (DEL-1) expression promotes an immunoprotective phenotype in experimental visceral leishmaniasis

**DOI:** 10.1101/2024.04.24.587740

**Authors:** Jakub Lukaszonek, Najmeeyah Brown, Jessie Zenga, Elmarie Myburgh, James P. Hewitson, George Hajishengalis, Triantafyllos Chavakis, Paul M. Kaye, Ioannis Kourtzelis

**Affiliations:** Hull York Medical School, University of York, United Kingdom; York Biomedical Research Institute, University of York, United Kingdom; Department of Biology, University of York, United Kingdom; Department of Basic and Translational Sciences, Penn Dental Medicine, University of Pennsylvania, Philadelphia, PA, USA; Institute for Clinical Chemistry and Laboratory Medicine, University Hospital and Faculty of Medicine, Technische Universität Dresden, Dresden, Germany

## Abstract

The identification of tissue-derived homeostatic molecules regulating immune plasticity is essential for understanding the role of macrophages in immune responses to intra-phagosomal pathogens. Developmental endothelial locus-1 (DEL-1) is a functionally versatile homeostatic factor capable of inhibiting the onset of inflammation and promoting inflammation resolution, but its role in the response to intracellular infections has not been previously addressed. *Leishmania*, causative agents of the neglected tropical disease leishmaniasis, are intra-phagosomal parasites that establish a replicative niche within macrophages.

Here, using a well-established murine model of visceral infection with *Leishmania donovani*, we establish DEL-1 as a novel regulator of immunity to this infection. Parasite burden was significantly higher in B6.*Edil3*^-/-^ (Del1-KO) compared to wild type B6 mice, as determined by whole body IVIS imaging, largely as a result of increased liver parasite load. However, lack of DEL-1 enhanced hepatomegaly and enhanced granulomatous inflammation. Conversely, parasite burden and the formation of large granulomas was reduced in mice overexpressing DEL-1 in macrophages but not in endothelial cells. Our findings reveal a hitherto unknown role of DEL-1 in the immune response to *L. donovani* infection and may represent a novel approach to mitigate immunopathology.

## Introduction

Macrophages are resident in tissues or can be differentiated from monocytes that are recruited from the periphery into the inflamed sites^1^. Both monocyte-derived and resident macrophages can detect, engulf and eliminate intra-phagosomal pathogens, such as mycobacteria, *Leishmania* and *Salmonella*^2^. However, macrophages in liver, spleen and bone marrow can also act as a safe haven for such pathogens, allowing them to reside and multiply, depending on their differentiation status^2,3^. Failure to eliminate these pathogens can cause macrophages to release large amounts of pro-inflammatory molecules that promote potentially damaging inflammation^3-5^. Thus, deciphering the multi-level interactions between macrophages, intra-phagosomal pathogens and other pathways that regulated inflammation may offer considerable therapeutic potential.

We have previously shown that the secreted protein developmental endothelial locus-1 (DEL-1) interacts with β2 and β3 integrins to modulate both the initiation and resolution of sterile inflammatory responses^6^. Specifically, DEL-1 (also known as epidermal growth factor (EGF)-like repeats and discoidin-I-like domains 3; EDIL3) is an endogenous negative regulator of inflammation expressed by several immune and non-immune cell types including macrophages and endothelial cells^6,7^. This protein controls β2 integrin-dependent leukocyte adhesion by blocking the interaction between the leukocyte integrin αLβ2 (LFA-1) and its endothelial counter-receptor ICAM-1^8^, as well as the binding of the αMβ2 (Mac-1) integrin with its ligand complement fragment iC3b^9^. On the basis of its ability to control β2 integrin activities, DEL-1 has emerged as a regulator of acute and chronic inflammatory responses in several disease models, including lung inflammation, neuroinflammation and inflammatory bone loss^6-8,10-13^. We have also demonstrated that DEL-1 functions as a non-redundant downstream effector in clearance of non-infectious inflammation^6^. Along this line, DEL-1 enhances the uptake of apoptotic neutrophils by interacting with the integrin receptor αvβ3 thereby promoting macrophage reprogramming into a pro-resolving phenotype in sterile inflammatory models^6,7,14^. While the role of DEL-1 in promoting resolution of non-infectious inflammatory responses is established, whether DEL-1 acts as an immune modulator to shape host responses to infections triggered by intra-phagosomal pathogens remains poorly explored.

*Leishmania spp*. are protozoan parasites that cause leishmaniasis, a collective of neglected tropical diseases with highly variable clinical forms that affect millions of people worldwide^15,16^. *Leishmania donovani* infection is associated with visceral leishmaniasis (VL) a systemic disease that is usually fatal if untreated^15,16^. Current drugs have significant limitations with the potential for further emergence of drug resistance^15,17,18^. No vaccines are available for use in humans^15^. Experimental rodent models of VL are well established^5,8^ and although not mimicking the full breadth of clinical disease, they represent an important tool for addressing mechanisms of host protection and / or immunopathology. Here, we studied the role of DEL-1 in host resistance and the immunopathology associated with systemic infection with *L. donovani*. Our data demonstrate that macrophage-derived DEL-1 plays a hitherto unrecognized role in the control of hepatic parasitic burden and in regulating the extent of granulomatous inflammation. This role for DEL-1 in regulating response to intra-phagosomal pathogens has the potential to be harnessed therapeutically.

## Results

### Parasitic burden is increased in mice deficient in DEL-1

Having previously identified DEL-1 as an endogenous negative immune regulator in the course of sterile inflammatory responses^6^, we sought to determine its role as a regulator of immunity during experimental VL following infection with a well-characterised luciferase-expressing line of *L. donovani*^19^ (**Fig. 1A)**. Following intravenous injection of *L. donovani* amastigotes, total body parasite load determined by IVIS imaging was equivalent 7 days post infection (p.i.) in wild type B6 mice and mice lacking DEL-1 (Del1-KO), suggesting that DEL-1 played only a minor role in the early stages of infection. In contrast, whereas parasite burden increased 4-fold over the subsequent 3 weeks in B6 mice, it increased ∼8-fold in Del1-KO mice **(Fig. 1B, C)**. As IVIS imaging does not clearly distinguish between parasites in spleen and liver, we directly analysed tissue parasite loads by the impression smear technique. At day 28 p.i., parasite load in the liver (**Fig. 1D**) but not spleen (**Fig. 1E**) was significantly greater in Del1KO mice compared to B6 mice, suggesting that liver parasite burden likely accounted for most of the differences observed using IVIS imaging. *L. donovani* infection in mice is characterised by enlargement of spleen and liver. Hepatomegaly is linked to granulomatous inflammation, a response that focuses anti-parasite effector responses on infected Kupffer cells and prevents collateral hepatic damage. In agreement with the elevated parasite load in the liver of Del1-KO mice, hepatomegaly (**Fig. 1F**), but not splenomegaly (**Fig. 1G**) was also increased in these mice. Histological analysis of liver sections stained with haematoxylin and eosin (H&E) revealed that Del1-KO mice had a significantly greater proportion **(Fig. 1H)** and abundance **(Fig. 1I)** of larger granulomas compared to WT mice.

**Figure 1:**
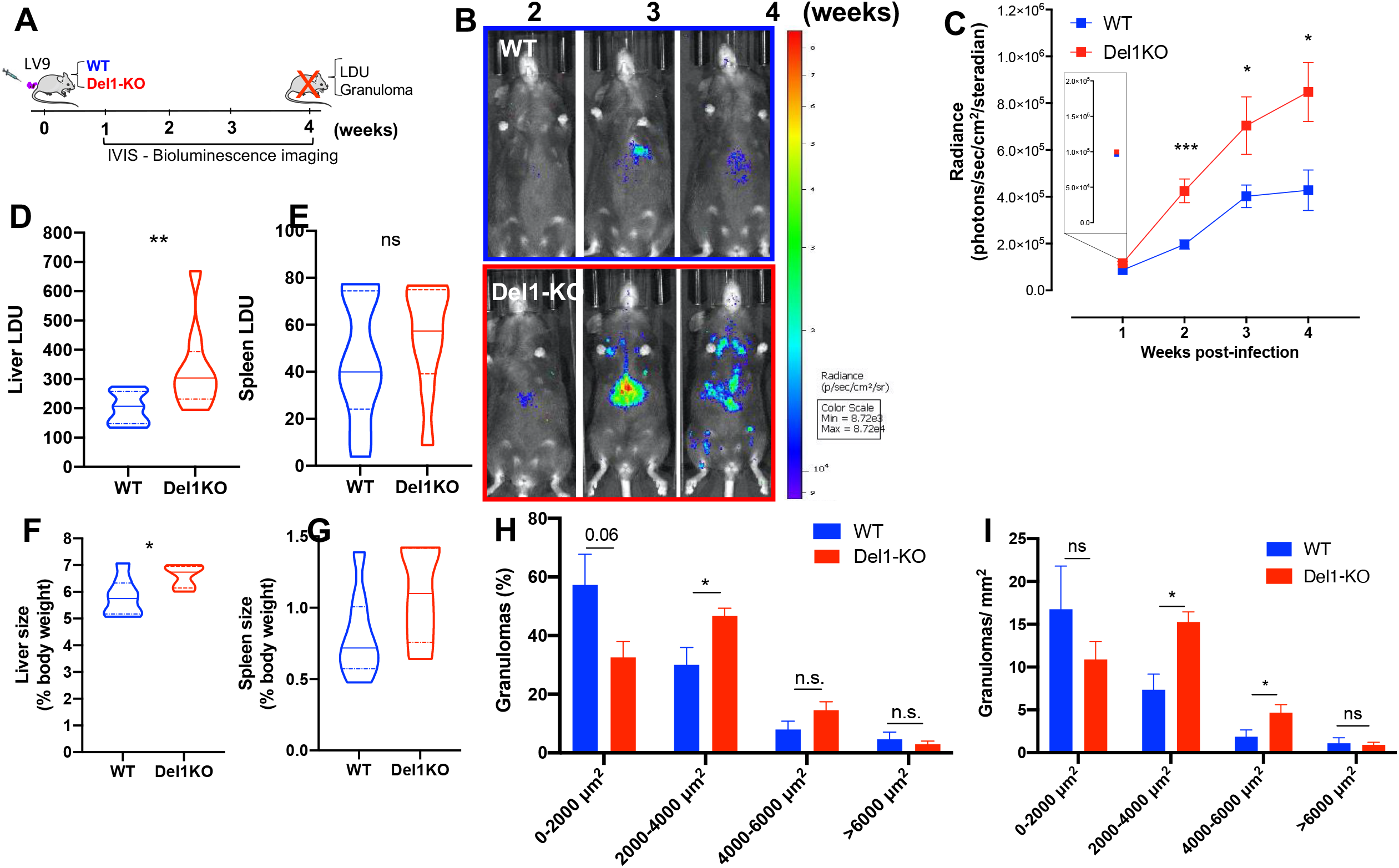
DEL-1 deficiency contributes to spread of *L. donovani*. **A**. Schematic diagram of experimental layout. **B, C**. Mice were infected with 3×10^7^ amastigotes of the bioluminescent luciferase-expressing *L. donovani LV9-*RE9H and imaged ventrally with IVIS bioluminescence *in vivo* imaging weekly after infection to assess the parasite loads in tissues. *In vivo* detection of *L. donovani* parasites in wild type (WT) and Del1-KO mice. Bioluminescence signal was converted to radiance (photons/sec/cm^2^/steradian; n = 9 mice / group, two-way ANOVA with Sidak’s multiple comparisons test for comparisons with corresponding time point). **D, E**. Day 28 parasite burdens expressed as LDU (Leishman-Donovan units; n = 9 mice / group, two-tailed Mann-Whitney *U*-test) in liver (D) and spleen (E). **F, G**. Liver (**F**) and spleen (**G**) size expressed as % body weight for infected mice. **H, I**. Size and numbers of hepatic granulomas were determined with H&E staining (n = 6 mice / group). (**A-G, I**) Data are presented as mean ± s.e.m or median with quartiles \**P* < 0.05, \*\**P* < 0.01, n.s., non-significant. (**H**) Percentage distribution of hepatic granulomas by size in infected mice (*P* = 0.005).

### DEL-1 deficiency enhances the pro-inflammatory phenotype in the infected liver

We next investigated the consequences of DEL-1 deficiency on immune cell phenotype and activity. Mice were infected with *L. donovani* and after four weeks immune cell subsets orchestrating the hepatic granulomatous response were analysed with flow cytometry. Neutrophils, monocytes, macrophages and lymphocytes were present in comparable proportions in the liver of infected Del1-KO and WT mice (**Supplementary Fig. 1**). IFNγ production by CD4^+^ T cells is cardinal feature of host resistance to *Leishmania* and its production is reflected in monocyte phenotype^20-25^. In uninfected control mice, the absence of DEL-1 had no effect on the proportion of CD4^+^ T cells secreting IFNγ (**Fig. 2A)** or the level of expression (**Fig. 2B**). Similarly, we observed no difference in inducible nitric oxide synthase (iNOS), major histocompatibility complex II (MHC-II) and TNF expression in Ly6C^high^ monocytes / macrophages (**Fig. 2C-E**). In contrast at day 28 p.i., the frequency of CD4^+^ T cells with commitment to IFNγ production was increased in Del1-KO mice compared to B6 mice (**Fig. 2F-H**) and this was reflected by an increase in iNOS (**Fig. 2I, K**), MHC-II; **Fig. 2L, M**) and TNF (**Fig. 2N, O**) expression by inflammatory Ly6C^high^ monocytes. Along the same line, cell numbers of IFNγ^+^ CD4^+^ T helper cells, iNOS^+^ Ly6Chigh and MHC-II^+^ Ly6Chigh monocytes / macrophages were increased in Del1-KO mice compared to B6 mice (**Supplementary Fig. 2**).

**Figure 2:**
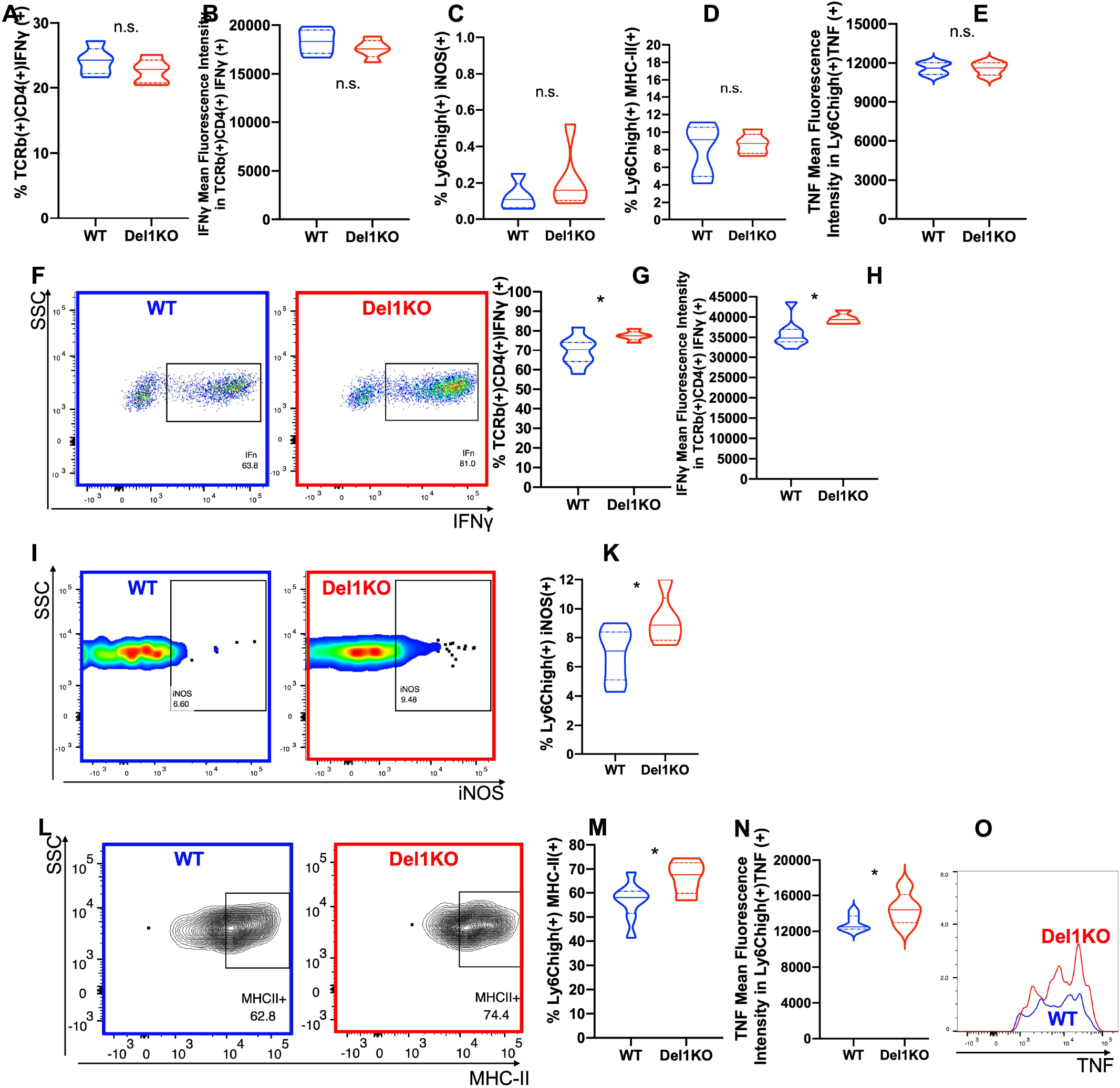
DEL-1 contributes to immune cell activity in infected liver. (**A**) Percentage and (**B**) mean fluorescence intensity (MFI) of IFNγ^+^ TCRβ^+^CD4^+^ cells from uninfected mice determined by intracellular cytokine staining following 4 h *ex vivo* stimulation with PMA and ionomycin. (**C**) Percentage of iNOS^+^ Ly6Chigh cells from uninfected mice determined by intracellular cytokine staining. (**D**) Percentage of MHC-II^+^ Ly6Chigh cells from uninfected mice. (**E**) MFI of TNF^+^ Ly6Chigh cells from uninfected mice, determined by intracellular cytokine staining following 4 h *ex vivo* stimulation with LPS. **F - H**. Representative FACS plot (**F**), percentage (**G**) and mean fluorescence intensity (**H**) of IFNγ^+^ TCRβ^+^CD4^+^ cells from d28 *L. donovani* - infected mice, determined by intracellular cytokine staining following 4 h *ex vivo* stimulation with PMA and ionomycin. **I, K**. Representative FACS plot (**I**) and percentage (**K**) of iNOS^+^ Ly6Chigh cells from d28 *L. donovani* - infected mice, determined by intracellular cytokine staining. **L, M**. Representative FACS plot (**L**) and percentage (**M**) of MHC-II^+^ Ly6Chigh cells from d28 *L. donovani* - infected mice. **N, O**. Mean fluorescence intensity (**N**) and representative FACS histogram (**O**) of TNF^+^ Ly6Chigh cells from d28 *L. donovani* - infected mice, determined by intracellular cytokine staining following 4 h *ex vivo* stimulation with LPS. Cells were gated as live intact singlets CD45^+^CD11b^-^CD8^-^CD19^-^TCRβ^+^CD4^+^ (**A, B, F - H**), CD45^+^SiglecF^-^Ly6g^-^CD64^+^CD11b^+^Ly6Chi^+^ (**C, D, I - M**), Cells were gated as live intact singlets CD45^+^SiglecF^-^Ly6g^-^F4/80^+^CD11b^+^Ly6Chi^+^TNF^+^ (**E, N, O**). Data are presented as median with quartiles, \**P* < 0.05 (two-tailed unpaired *t*-test; n.s., non-significant). A - E: n = 5 mice per group, F – O: n = 8 WT and 5 Del1-KO mice.

Therefore, in contrary to what might have been expected from the elevated parasite loads, infected Del1-KO mice show enhanced pro-inflammatory responses to infection.

### Over-expression of DEL-1 in macrophages enhances host resistance but blunts inflammation

To further understand why DEL-1 deficiency led to an elevated parasite burden despite an enhanced proinflammatory response (**Fig. 1 and 2**), we examined the consequences of DEL-1 over-expression. We have previously shown a cell type – dependent role of DEL-1 in acute self-limited sterile inflammation^6^. Specifically, macrophage-derived DEL-1 (generated using the human *CD68* promoter) promotes resolution of inflammation, whereas endothelial cell–derived DEL-1 regulates leukocyte recruitment. Hence, we infected mice with endothelial-specific overexpression of DEL-1 (EC-Del1)^10^ and mice with macrophage-specific DEL-1 overexpression (MΦ-Del1)^6^ (**Fig. 3A**). DEL-1 levels in these transgenic lines have previously been demonstrated^6,10^. EC-Del1 mice had total body parasite loads comparable to WT mice as determined by IVIS imaging (**Fig. 3B**). In contrast, parasite burden was decreased in MΦ-Del1 compared with WT infected mice (**Fig. 3B, C**). Similarly, at 10 weeks post-infection, liver parasite burden determined microscopically was lower in MΦ-Del1 compared to WT mice (**Fig. 3D**). DEL-1 over-expression in macrophages or endothelial cells did not affect parasite load in spleen (**Fig. 3E**), in keeping with the lack of an effect of DEL-1 KO in this organ (**Fig. 1**). Enhanced resistance in MΦ-Del1 mice was accompanied by a reduction in the proportion of large hepatic granulomas compared to WT and EC-Del1 mice (**Fig. 3F, G**). These findings suggest that endothelial cell-derived DEL-1, unlike previously shown in sterile inflammatory models^6,7,10^, may not modulate inflammatory outcome in experimental visceral leishmaniasis. In contrast, DEL-1 overexpression in macrophages leads to enhanced resistance and dampening of inflammation.

**Figure 3:**
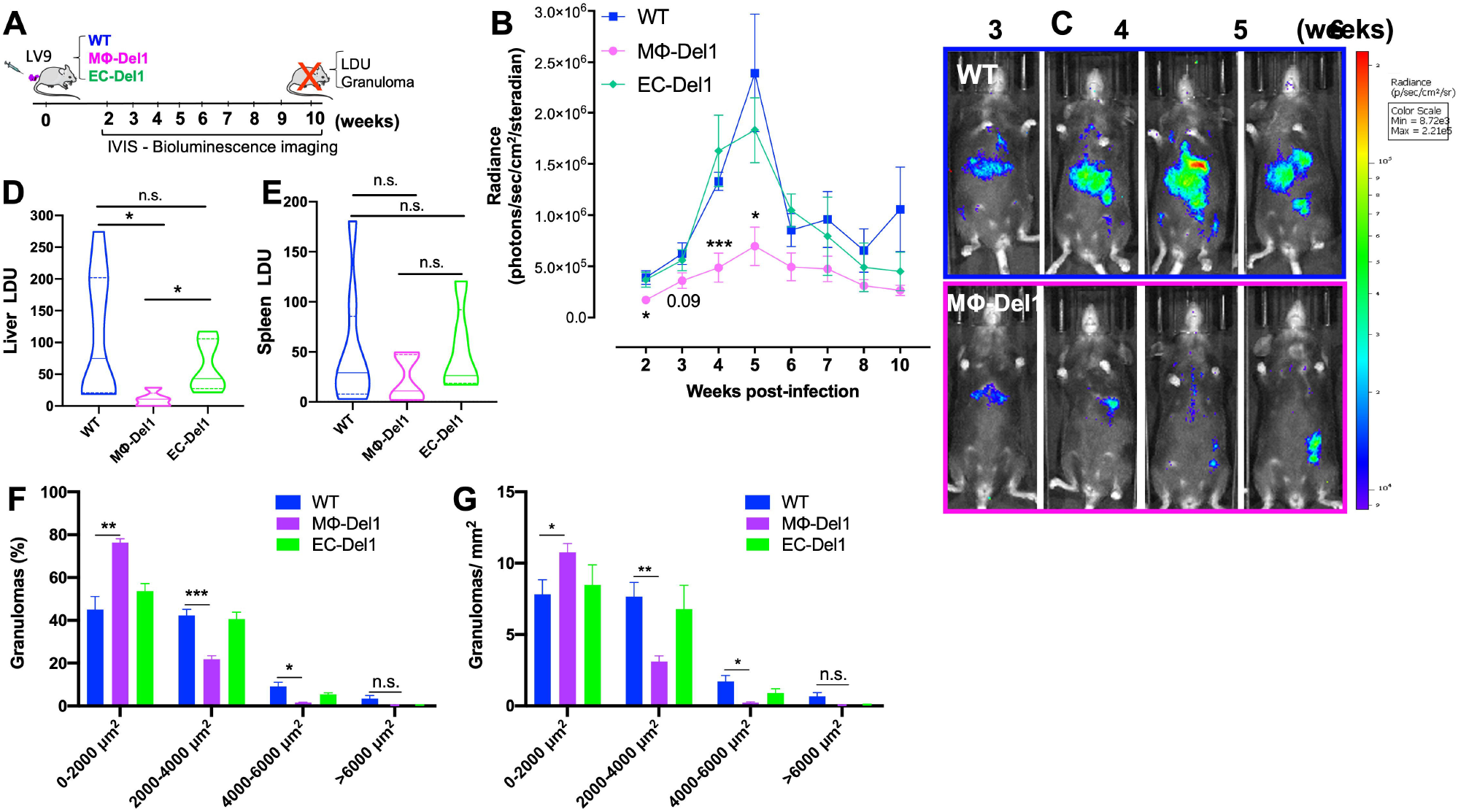
Cellular sources of DEL-1 differentially regulate L. *donovani* burden. **A**. Schematic diagram of experimental layout. **B, C**. Mice were infected with 3 × 10^7^ amastigotes of the bioluminescent luciferase-expressing *L. donovani LV9-*RE9H and imaged ventrally with IVIS bioluminescent *in vivo* imaging after infection to assess the parasite loads in tissues. *In vivo* detection of *L. donovani* parasites in wild type (WT), endothelial-specific DEL-1 overexpressing (EC-Del1) and macrophage-specific DEL-1 (MΦ-Del1) overexpressing mice. Bioluminescence signal was converted to radiance (photons/sec/cm^2^/steradian; two-way ANOVA with Sidak’s multiple comparisons test for comparisons with corresponding time point). **D, E**. Week 10 parasite burdens expressed as LDU (Leishman-Donovan units; (one-way ANOVA with Tukey’s multiple comparisons test) in liver (D) and spleen (E). **F, G**. Size and numbers of hepatic granulomas were determined at week 10 with H&E staining. (**A-E, G**) Data are presented as mean ± s.e.m. or median with quartiles, n = 8 WT, 5 MΦ-Del1, 5 EC-Del1 mice. \**P* < 0.05, \*\**P* < 0.01, \*\*\**P* < 0.001; n.s., non-significant. (**F**) Percentage distribution of hepatic granulomas by size in infected mice (*P* < 0.0001).

Together, these data unveil a novel role for DEL-1 in shaping the immune response to L. *donovani* infection, providing a better understanding on mechanisms that may shift the balance of damaging inflammation towards pathogen elimination.

## Discussion

Host defence mechanisms against *L. donovani* infection involves several immune cell types and inflammatory events including CD4^+^ T helper cell activity, monocyte infiltration, macrophage activation and cytokine release as well as granuloma formation in liver, all of which aim to contribute to parasite elimination^20-22,24,27^. Although immune cell interactions play important roles in the immunopathology of leishmaniasis, the network of molecules and mechanisms that coordinate this complex process is incompletely understood. Therefore, identification of endogenous immune regulators that aim to attenuate parasite burden is required to harness chronic destructive inflammatory responses.

In sterile inflammatory disease models, the endogenous negative regulator of inflammation DEL-1 inhibits leukocyte adhesion^6,11^ and promotes inflammation resolution^6^. Here we investigated whether DEL-1 regulates immune responses during *L. donovani* infection. We show that DEL-1 deficiency limits host control of hepatic parasite burden and that is accompanied by the generation of larger hepatic granulomas. We also found that key inflammatory mediators such as IFNg, iNOS and TNF, all previously linked to enhanced anti-parasite immunity^21,23,24,27,29^, were up-regulated by immune cells infiltrated in the liver of infected Del1-KO mice. Further studies would be useful to address why Del1-KO mice do not control parasite burden, despite their ability to promote anti-*Leishmania* immune responses.

The role of DEL-1 on promoting host resistance to this intracellular pathogen was corroborated by showing that overexpression of DEL-1 in macrophages leads to decreased parasitic burden whereas mice overexpressing DEL-1 in endothelial cells had no detectable effect. The lower parasite load in infected MΦ-DEL1 mice was accompanied by blunting of the granulomatous response. Future studies using mice with a conditional Del-1 allele and a macrophage – specific deletion of DEL-1 could address in more detail the physiological relevance of macrophage – derived DEL-1 in this disease model. Parasite recognition and clearance by macrophages can lead to decreased parasitic burden^30-32^. Therefore, the action of DEL-1 on this model may be attributed at least in part to the modulation of pathogen phagocytosis, but further analyses are required to test this hypothesis.

Future studies should focus on the testing of the immunomodulatory role of DEL-1 using a pharmacological approach. Treatment of infected WT mice with different DEL-1 variants could facilitate drug development. In particular, use of DEL-1 protein either lacking the two discoidin I-like domains or a point mutant that has a single amino acid mutation at the RGD motif located in the second EGF-repeat^12^ could allow the identification of structural components of DEL-1 involved in the anti-parasitic responses. Importantly, DEL-1 has already been used therapeutically to modify osteoclastogenesis in non-human primates^12^. Intriguingly, both the macrolide antibiotic erythromycin and the non-antibiotic immunomodulatory derivative^33,34^ upregulate DEL-1 expression providing an inexpensive regimen to modulate DEL1 therapeutically.

In summary, the present study pinpoints a novel role for DEL-1 in attenuating hepatic parasite burden during experimental visceral leishmaniasis and demonstrates that the anti-parasitic effects of DEL-1 are dependent on cell-specific expression. These findings may apply to other diseases caused by pathogens with an intra-phagosomal lifestyle.

## Materials and Methods

### Mice and parasite infections

#### Mice

C57BL/6 (B6) mice, Del1 KO mice, EC-Del1 mice (overexpressing DEL-1 in the endothelium) and MΦ-Del1 mice (overexpressing DEL-1 in macrophages)^6,10^ were bred in-house under specific pathogen-free conditions on a standard 12/12 h light/dark cycle according to the institutional guidelines at the Biological Services Facility (BSF), University of York. Female mice that were 8-10 weeks old were used in experimental procedures. Food and water were provided ad libitum.

#### Parasite infections

The Ethiopian strain of *L. donovani* (LV9) and bioluminescent *LV9-*RE9H^19^ parasites were maintained by passage in female *Rag2*-KO mice. DEL-1 deficient mice, transgenic lines with cell-specific overexpression of DEL-1, littermate controls and WT mice were infected intravenously (i.v.) with 3 × 10^7^ amastigotes *via* lateral tail vein. Use of littermate controls showed similar results to those obtained from WT mice (data not shown). All animals were visually inspected daily and were within accepted humane endpoints.

### *In vivo* bioluminescence imaging

IVIS Spectrum *in vivo* imaging system (PerkinElmer, UK) was used to monitor luminescence in mice infected with the bioluminescent *LV9-*RE9H parasites^26^. Mice were injected intraperitoneally with D-Luciferin potassium salt (150 mg / kg; Syd Labs, USA) and after 5 min they were anaesthetized with gaseous isoflurane. Anaesthetised mice were kept under anaesthesia and were imaged 10 minutes after D-Luciferin administration in a group of 3 animals for 4 minutes on ventral side. Emission filter was open for 30-60 seconds exposure, large binning and 1 f/stop. After imaging, mice were monitored and kept on a heating pad until fully recovered from the anaesthesia. Image analysis was performed using Living Image Software (PerkinElmer, UK) by drawing regions of interest of equal size around the whole body to quantify bioluminescence. Bioluminescence signal was converted to radiance (photons/sec/cm^2^/steradian) using a fixed normalised scale on IVIS images of animals.

### Tissue sectioning and histology

Liver tissues from infected mice were harvested at the end of each experiment, were embedded in cryomolds using OCT Tissue-Tek (Labtech) and snap frozen on dry ice. Cryo-embedded tissue sections with thickness 8 μm were allowed to air dry prior to haematoxylin and eosin staining. Coverslips with DePeX mounting medium were then added to the slides and images were captured using an AxioScan.Z1 slide scanner (Zeiss) at 20x resolution using Zen software (Zeiss). Granuloma size was defined by area. Data were collected from 1–3 sections per mouse and pooled for image analysis using ImageJ software.

### Determination of parasite burden

*Leishmania* parasite burden was calculated based on counts from Giemsa-stained tissue impression smears. It is expressed as Leishman-Donovan units (LDU), where LDU is equal to the number of parasites per 1000 host cell nuclei multiplied by the organ weight in milligrams.

### Leukocyte isolation from liver

Liver single cell suspensions were prepared after digestion with liberase TL (0.8 U/ml; Roche) and DNase (160 U/ml; Sigma) in Hank’s Balanced Salt Solution for 45 min at 37°C with shaking. Enzyme activity was inhibited with 10 mM EDTA pH 7,5 and preparations were passed through a 100-μm nylon cell strainer (BD Biosciences) in complete RPMI 1640 (ThermoFisher) supplemented with 10% heat-inactivated fetal calf serum, 100 U/ml penicillin, 100 μg/ml streptomycin and 2 mM L-glutamine (ThermoFisher). Leukocyte isolation was performed using Percoll gradient followed by red blood cell lysis (Sigma).

### Flow cytometric analysis

Live/dead cell discrimination was performed using zombie aqua (BioLegend) and subsequently Fc receptors were blocked with rat IgG (100 μg/ml; Sigma). Cell surface staining was performed in FACS buffer (PBS / 0.5% BSA / 0.05% azide) for 30 min at 4°C. For phenotypic analysis, leukocytes were stained with combinations of the following anti-mouse antibodies: CD45 BV785 (104), TCRβ PE-Cy7 (H57-597), CD4 PerCP/Cy5.5 (RM4-5); CD8α APC (53-6.7), Siglec-F e710 (1RNM44N), Ly6G APC-Cy7 (1A8), CD11b PB and PE-Cy7 (M1/70), Ly6C BV605 (HK1.4), CD64 PE (X54-5/7.1), MHCII BV650 (M5/114.15.2), F4/80 FITC (BM8), CD19 APC-Cy7 (6D5), IFNγ FITC (XMG1.2), iNOS e610 (CxNF7), IL-4 PE-dazzle (11B11), IL-10 PE (JES5-16E3), TNF BV421 (MP6-XT22). Antibody against Siglec-F was from eBiosciences and all other antibodies were from BioLegend.

To measure intracellular cytokines in T cells, leukocytes were stimulated *ex vivo* with 500 ng/ml Phorbol-12-myristate-13-acetate (PMA; Sigma-Aldrich) and 1 μg/ml ionomycin (Sigma-Aldrich) in the presence of 10 μg/ml brefeldin A (Sigma) for 4 h at 37°C. To measure intracellular cytokines in myeloid cells, cells were cultured in the presence of brefeldin A and LPS (1 μg/ml *Escherichia coli* O55:B5; Sigma). For intracellular cytokine staining, surface-stained cells were either treated with the Foxp3 Transcription Factor Fixation / permeabilisation buffer (for iNOS, eBiosciences) or IC Fixation/permeabilisation buffer (for intracellular cytokines, BD). Negative controls were stained with matched-isotype controls or FMO (fluorescent minus one) samples. Flow cytometric analysis was performed on a LSR Fortessa (BD Biosciences) and data were analysed with FlowJo software (TreeStar).

### Statistical analysis

Statistical analyses were carried out with GraphPad Prism software. For infection experiments, researchers were blinded to mouse genotype. This also applies to histology and flow cytometry analyses. Results are presented as mean plus standard error of the mean or median with quartiles. Two tailed parametric and non-parametric tests were used as appropriate after testing for normality. Multiple-group comparisons were performed using analysis of variance (ANOVA) and the Tukey’s or Sidak’s multiple comparison tests. Distribution data were analysed by chi-squared test. A *P* value of < 0.05 was considered statistically significant. All statistical information is provided in each figure legend.

### Ethics statement

All animal experiments were carried out under the authority of a UK Home Office Licence (project licence number PPL PP0326977) that obtained approval by the University of York Animal Welfare and Ethics Review Board. All procedures were performed in compliance with ARRIVE guidelines. No unexpected adverse events were recorded during this study.

## Author contributions

JL and JZ performed experiments, analysed and interpreted data; NB designed and performed experiments, analyzed and interpreted data; EM generated reagents; JPH, GH, TC contributed to experimental design and interpreted data; PMK contributed to experimental design, interpreted data and supervised research; IK conceived and designed the study, performed experiments, supervised research, interpreted data, and wrote the paper.

All authors critiqued and edited the manuscript.

## Acknowledgements

We would like to thank staff at the Imaging and Cytometry Lab in the University of York Bioscience Technology Facility for imaging support and advice. We thank Bruce Branchini (Connecticut College, US) for the PpyRE9H red shifted luciferase gene used the generate LV9-RE9H.

---

**Supplementary Figure 1: Immune cell frequencies in infected liver**.

Percentage of neutrophils (**A**; gated as live intact singlets CD45^+^SiglecF^-^CD11b^+^Ly6g^+^), inflammatory monocytes / macrophages (**B**; gated as live intact singlets CD45^+^SiglecF^-^Ly6g^-^ CD64^+^CD11b^+^Ly6Chigh), macrophages (**C**; gated as live intact singlets CD45^+^SiglecF^-^Ly6g^-^ CD64^+^CD11b^+^Ly6Clow), CD4^+^ T cells (**D**; gated as live intact singlets CD45^+^CD11b^-^CD8^-^ CD19^-^TCRβ^+^CD4^+^), CD8^+^ T cells (**E**; gated as live intact singlets CD45^+^CD11b^-^CD4^-^CD19^-^ TCRβ^+^CD8^+^), B Cells (**F**; gated as live intact singlets CD45^+^CD11b^-^TCRβ^-^CD19^+^) from day 28 *L. donovani* - infected mice. Data are presented as median with quartiles, n = 8 WT and 5 Del1-KO mice (n.s., non-significant, two-tailed unpaired *t*-test).

**Supplementary Figure 2: Cellularity of immune cell types in infected liver**.

Number of CD4^+^ T cells (**A**), IFNγ^+^ CD4+ T cells (**B**), iNOS^+^ Ly6Chigh monocytes (**C**), MHC-II^+^ Ly6Chigh monocytes (**D**) and monocytes (**E**) from d28 *L. donovani* - infected mice. Cells were gated as described in **Fig. 2**. Data are presented as median with quartiles, \**P* < 0.05 (two-tailed unpaired *t*-test; n.s., non-significant, n = 8 WT and 5 Del1-KO mice).

