## Supplementary figures and images for "Macrophage-derived developmental endothelial locus 1 (DEL-1) expression promotes an immunoprotective phenotype in experimental visceral leishmaniasis"

### Supplemental Figure 1

#### Supplementary Figure 1

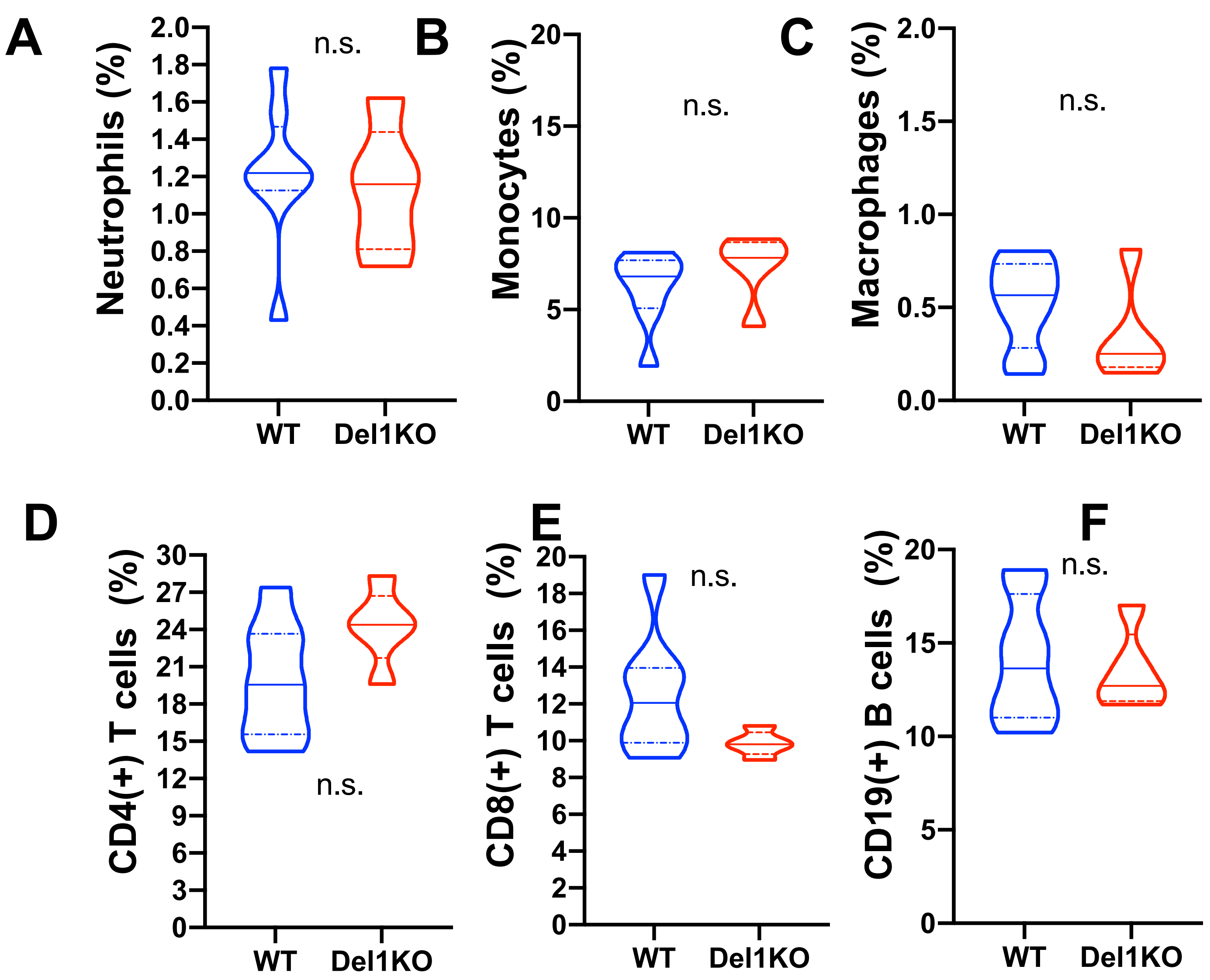

### Supplemental Figure 2

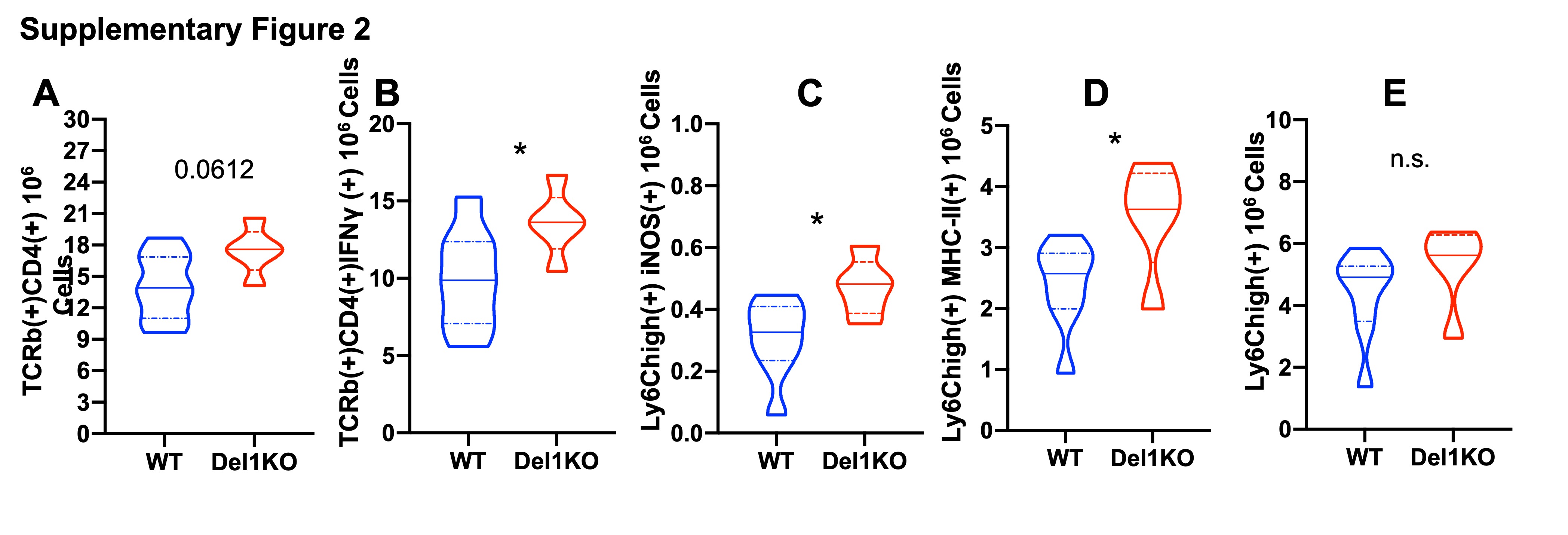
